# Propagation of Beta bursts from the motor cortex to the motor units of multiple upper-limb muscles

**DOI:** 10.1101/2025.05.22.655541

**Authors:** Cosima Graef, Alejandro Pascual Valdunciel, Dario Farina, Ravi Vaidyanathan, Yen Tai, Shlomi Haar

## Abstract

Beta band (13-30 Hz) oscillations are crucial for motor control, though their functional significance remains debated. Recent research suggests that beta activity occurs in transient bursts, which may better capture its role in movement regulation than sustained oscillations. While cortical and sub- cortical beta bursts have been extensively studied, their transmission to muscles—particularly in the upper limb—remains poorly understood and has been limited by traditional bipolar EMG techniques. In this study, we used high-density surface electromyography (HDsEMG) and electroencephalography (EEG) to investigate the cortico-peripheral dynamics of beta bursts in forearm extensor muscles during isometric contractions at the motor unit (MU) level. We show that MU activity in the upper limb exhibits discrete beta bursts that are temporally aligned with cortical beta activity. Notably, beta bursts in the periphery were time-locked to cortical bursts, suggesting strong coordination and synchronisation of bursting across the corticospinal tract. We also found stronger beta synchronisation in the extensor carpi ulnaris compared to the extensor carpi radialis, indicating muscle-specific differences in shared neural drive. These findings provide the first demonstration of beta burst propagation from cortex to upper-limb MUs and show that HDsEMG can reliability detect such events in the upper limb. This work supports the cortical origin and structure of peripheral beta activity and demonstrates its potential as a neurophysiological biomarker for targeting corticospinal dynamics in motor disorders such as Parkinson’s disease.

**Significance Statement:** We provide the first evidence of cortical beta bursting in motor unit (MU) activity of forearm extensors across varying force levels. Our results show that MU beta bursting is time-locked to cortical beta bursts, supporting the idea of direct corticospinal transmission in the upper limb. Our findings highlight the value of MU level analysis in understanding beta burst transmission and indicate that burst timing is robust to changes in contraction levels. Unlike traditional bipolar EMG, MU-level analysis offers higher temporal precision and source specificity, enabling the reliable detection of beta bursts in individual muscles. These findings demonstrate a robust cortical-peripheral beta relationship during isometric contractions and reinforces the utility of HDsEMG for investigating burst dynamics and refining neurophysiological biomarkers.

## Introduction

Oscillatory activity in the beta frequency range (13–30 Hz) plays a fundamental role in motor control. It has traditionally been thought to stabilize and maintain consistent motor output (Engel & Fries, 2010), while its synchrony between the motor cortex and muscles supports the fine-tuning of ongoing movements (Kilavik et al., 2013). However, recent evidence suggests that beta activity is not sustained continuously but instead occurs in brief, transient bursts (Feingold et al., 2015). These bursts encode functionally relevant information for movement initiation (Little et al., 2019) and performance (Torrecillos et al., 2018), and are implicated in motor disorders such as Parkinson’s disease (PD) (Tinkhauser, Pogosyan, Little, et al., 2017; Tinkhauser, Pogosyan, Tan, et al., 2017a). Although most research on pathological beta activity has focused on the subthalamic nucleus (STN) (Tinkhauser, Pogosyan, Tan, et al., 2017b), cortical beta bursts also play a significant role in motor processing (O’Keeffe et al., 2020; Pauls et al., 2022; Vinding et al., 2020). Notably, treatments like levodopa and deep brain stimulation (DBS) can reduce beta burst duration and rate, with beta-sensitive closed-loop DBS approaches being explored at both cortical and STN levels (Abbasi et al., 2018; Lofredi et al., 2019; Luoma et al., 2018; Spooner et al., 2024; Tinkhauser, Pogosyan, Little, et al., 2017; Tinkhauser, Pogosyan, Tan, et al., 2017a). A recent study found that cortical bursts precede STN bursts and better relate to worsening bradykinesia (Yao et al., 2025), highlighting the importance of tracking beta burst propagation.

Cortical beta activity propagates to the periphery via corticospinal pathways and its coupling with contralateral muscle activity during steady isometric contraction is evident in the corticomuscular coherence (CMC), which peaks in the beta frequency range (Brovelli et al., 2004; Salenius et al., 1997). Beta oscillations have been shown to transmit linearly from the motor cortex to tonically active muscles (Ibáñez et al., 2021). Both descending corticospinal pathways and ascending sensory inputs contribute to CMC, with coherence strength modulated by movement and muscle activity (Witham et al., 2011a). High-density electromyography (HDsEMG) enables detection of beta bursts in individual motor units (MUs), providing a direct measure of corticospinal beta transmission (Bräcklein et al., 2022). This offers a novel opportunity to bridge the gap in our understanding of how beta bursts propagate through the corticospinal tract and influence MU activity. Additionally, investigating beta bursts across multiple force levels may provide novel insights into how MUs synchronise with beta oscillations and how corticospinal transmission adapts to changes in force output, potentially informing models of motor impairment in PD.

To date, only two studies have tracked the propagation of beta bursts from the cortex to the MUs, both focusing on the tibialis anterior in the lower limb (Abbagnano et al., 2025; Bräcklein et al., 2022). Other studies that tracked beta propagation have relied on bipolar EMG recordings to examine CMC (Echeverria-Altuna et al., 2022; Simpson et al., 2024), but these lack the spatial resolution to isolate individual MU activity. The extent to which beta bursts exhibit similar transmission dynamics in the upper limb remains unclear, especially given the distinct corticospinal organization of upper limb muscles (Tinkhauser et al., 2019).

In this study, we investigate the cortical-peripheral relationship of beta bursts using EEG and HDsEMG during isometric forearm contractions. Specifically, we aim to: (1) assess the reliability of HDsEMG in detecting beta bursting in the upper limb, (2) determine whether peripherally recorded MU activity reflects cortical beta bursts, and (3) examine how beta bursts differ between the brain and muscle across different force levels. We hypothesize that beta bursts in forearm muscles will not exhibit distinct characteristics depending on contraction intensity given prior evidence that corticomuscular coherence across force levels stayed unchanged (Abbagnano et al., 2025). Additionally, we expect that cortical beta bursts will be time-locked with MU activity in the forearm extensors, supporting direct corticospinal transmission. By addressing these questions, this study seeks to advance understanding of the role of beta bursts in motor control and their potential implications for neurorehabilitation and closed-loop DBS strategies in PD.

## Materials and Methods

### Participants

We recruited 24 participants (26.0 ± 3.3 years, 6 females) who were all right-handed as self-reported. None of the participants had a reported history of neurological conditions or upper limb injuries. All participants gave written informed consent prior to participation in the study. The study was approved by the Imperial College Research Ethics Committee (ICREC number 6715837) and conducted in accordance with the Declaration of Helsinki.

### Experimental Design

At the start of the experiment, participants performed a single maximum voluntary wrist extension for 5 seconds to estimate the level of maximum voluntary contraction (MVC). To achieve an accurate maximum force level, verbal encouragement was provided. The MVC was calculated as the 95% percentile of the peak force value and was digitally recorded. This value was then used as a reference for the rest of the experiment to adjust the torque of the isometric contraction ramps to the participants’ individual strength levels.

Participants then familiarised themselves with the force trajectory task through practice trials at 10%, 20% and 30% MVC, where force trajectories and a trapezoidal force template were displayed on a monitor. Each trial consisted of 3s rest (baseline), a linear increase in force to the target contraction level for 4s (ramp up), 60s steady contraction at the target MVC level (plateau), 4s linear decrease in force to rest (ramp down), and 3s rest (baseline). The main experiment started after 3 min of rest and consisted of 3 blocks (1 at each MVC level: 10%, 20% and 30% MVC) with 5 trials per block. The order of MVC blocks was randomised, and there was a 60 second rest between 10% and 20% MVC trials and a 120 second rest between 30% MVC trials to minimise muscle fatigue effects.

### Data acquisition

#### Force recordings

The HRX-1 interface robot (HumanRobotix London, n.d.) was used to record torque during isometric wrist extension. This device was table-mounted and the participants’ dominant forearms were strapped to the device, with the back of the hand resting against the robot handle. Participants were seated at a height allowing the elbow joint angle to be set at 90° of flexion. The force signal was sampled at 24 Hz and recorded through custom MATLAB scripts. The force, and trapezoidal ramp visuals were displayed using the Psychophysics Toolbox Version 3 (Brainard, 1997; Kleiner et al., 2007; Pelli, 1997) written in MATLAB R2023a (The MathWorks Inc., 2023).

#### HDsEMG recordings

HDsEMG was acquired from the extensor carpi ulnaris and extensor carpi radialis muscles of the dominant forearm via a configuration of four 64-electrode grids (5 columns x 13 rows; gold-coated; 1- mm diameter; 4-mm interelectrode distance (IED); OT Biolettronica). Multiple grids were utilised in the configuration to maximise the total MUs identified from the HDsEMG recordings by using a large number of electrodes and short IED (Caillet et al., 2023). The grids were arranged to form a 256- electrode array (see figure 1A), with grids centred around the muscle belly whilst in parallel with the direction of muscle fibres. To improve signal quality, the skin was shaved, gently abraded using abrasive paste, and cleansed using a 70% ethyl alcohol solution. To maintain electrode-to-skin contact, a perforated bi-adhesive foam infused with conductive paste was used. The grids were fastened in place with tape and elastic bands. A wet band was positioned around the wrist of the same arm and used as reference to record the HDsEMG signals in monopolar mode. The HDsEMG signals were amplified (x150) using a multichannel EMG amplifier (Quattrocento, OT Bioelettronica, Turin, Italy), sampled at 2048 Hz by a 16-bit A/D converter, and bandpass filtered from 10 to 500 Hz. Signals were recorded with the OTBioLab software (OT Bioelettronica, n.d.).

**Figure 1.**
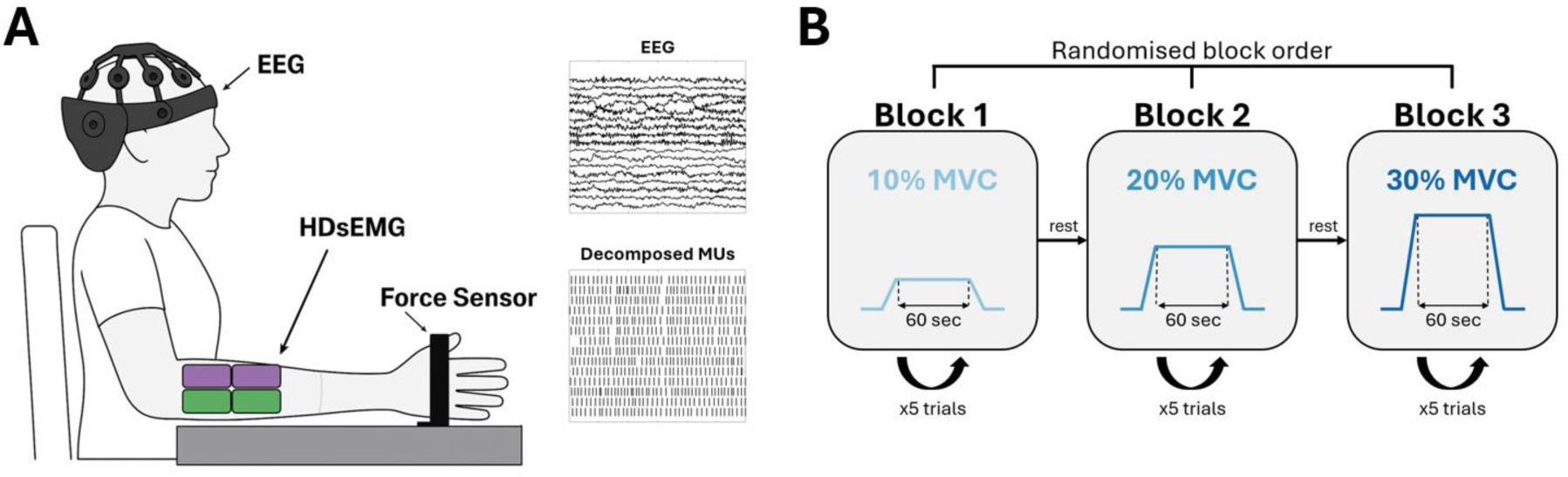
Overview of experimental setup and task paradigm. (A) Schematic of the EEG, HDsEMG, and torque recording setup during isometric wrist extension. Four HDsEMG grids were positioned over the extensor carpi ulnaris (ECU, green area) and extensor carpi radialis (ECR, purple area) muscles to record motor unit (MU) activity. EEG and HDsEMG signals were used to assess cortical-muscular interactions via cumulative spike trains (CSTs). (B) Participants performed a ramp-and-hold wrist extension task at 10%, 20%, and 30% maximum voluntary contraction (MVC) levels, following a randomized block design with rests in between trials and blocks.

#### EEG recordings

The EEG data was recorded synchronously with HDsEMG recordings, with 19 dry electrodes using the DSI-24 (Wearable Sensing, San Diego, CA, USA). The electrodes were arranged according to the International 10-20 system, Pz was used as the reference, Fpz as the ground, and signals were sampled at 300 Hz.

EEG, HDsEMG and force signals were synchronised using a hardware trigger connected between systems to ensure precise temporal alignment.

### Data Analysis

All offline analyses were conducted in MATLAB R2023a (The MathWorks Inc., 2023).

#### Decomposing HDsEMG into motor unit spike trains

The recorded HDsEMG monopolar signals were decomposed into underlying MU activity using a previously validated blind-source separation algorithm (Holobar et al., 2014; Negro et al., 2016), with each grid being decomposed separately. This decomposition algorithm’s accuracy has been demonstrated for isometric force contractions across a variety of muscles and force levels (Del Vecchio, Casolo, et al., 2019; Farina & Holobar, 2016; Holobar et al., 2009). Duplicate MUs within and between grids were identified and removed. MUs with more than 30% of the same discharge times were defined as duplicates. The decomposed discharge times of the MUs were manually inspected and edited in accordance with (Del Vecchio et al., 2020) to improve the decomposition accuracy using the MUedit software (Avrillon et al., 2024). The MU silhouette values (SIL) were calculated as a qualitative measure of evaluation of the decomposition accuracy. MUs with a SIL value of less than 0.9, firing delay of more than 5 seconds, or active window less than 50 seconds (of the 60 second plateau period) were discarded. Trials containing fewer than 5 MUs were excluded to ensure robust corticomuscular coupling estimates, based on prior findings that combining the activity of three or more MUs yields stable coherence and delay measurements (Ibáñez et al., 2021).

To allow a direct comparison of beta band activity across different MVC levels, we used a MU resampling approach to equalise the firing rates of cumulative spike trains (CSTs) across trials. This process made sure that variations in MU firing rate did not confound differences in beta activity, and that these differences were attributable to neural mechanisms. The minimum CST firing rate across all trials for each participant were determined, followed by iteratively selecting combinations of MUs within each trial to match this minimum firing rate, allowing for a 10% tolerance range. After MUs subsets were determined, new CSTs were reconstructed and coherence measures calculated. CSTs were obtained by summing binary MUs for the 60 second plateau period only. The CST average discharge rates were calculated by averaging the instantaneous discharge rates. For corticomuscular coherence (CMC) and bursting analysis, the MUs were downsampled to 300 Hz prior to calculating the CST.

#### EEG preprocessing

For the EEG preprocessing, the EEGLAB toolbox for MATLAB was used (Delorme & Makeig, 2004). The EEG signals were re-referenced to the linked ears, high pass filtered at 1Hz and then lowpass filtered at 45 Hz using a linear (zero-phase) non-causal FIR filter. This was done instead of bandpassing to avoid the problem of steeper than necessary low-pass slopes. Bad channels were removed automatically by removing flat channels (if flat for more than 5 s), channels with a substantial amount of noise (maximum acceptable high-frequency noise standard deviation of 4) and channels poorly correlated with other channels (minimum acceptable correlation with nearby channels of 0.8). Artifact subspace reconstruction was performed to correct small segments of bad data with a standard deviation cutoff for the burst removal of 6 and using a Euclidian distance. This was followed by performing independent component analysis (ICA) for artifact rejection. Components were flagged and removed if they had a more than 70% probability of being a muscle, eye or heart artifact. A current source density transform, more commonly known as a surface Laplacian, was applied to the EEG data using the CSD toolbox (Kayser & Tenke, 2006a, 2006b) as recommended by (Kayser & Tenke, 2015). EEG analysis was conducted on the C3 channel only due to its location over the sensorimotor cortex contralateral to the forearm according to the 10-20 system.

#### Coherence analysis

An essential aspect of our study involves examining the relationship between continuous EEG signals and spiking MU activity, as well as investigating intra-muscular coherence (IMC) of MUs from different extensor muscles and regions. The Neurospec 2.11 toolbox (David Halliday, 2016; Halliday, 2015) for MATLAB (The MathWorks Inc., 2023) which was previously validated (Beck et al., 2021; Ibáñez et al., 2021; Spedden et al., 2022), was integrated to calculate the coupling strength and frequency content between time series and point process data, or between 2 spike trains.

The IMC was estimated for trials with at least 2 MUs by splitting the MU pool into two randomly chosen groups of equal size and calculating their coherence using Neurospec’s two channel spectral analysis. The IMC was the average of coherence estimates over 100 iterations. The CMC was estimated between the downsampled to 300 Hz CST, and EEG C3 channel using Neurospec’s two channel spectral analysis. Both the IMC and CMC were estimated for MUs from individual grids, pairs of grids and all 4 grids together.

All coherence analysis was conducted with segments of 2 seconds and multitapers (three tapers). Significant coherences were identified using the upper 95% confidence limit estimates from Neurospec (Rosenberg et al., 1989). These are calculated using the formula 1−0.05^(1/(L−1)), where L represents the number of segments used in computing the Fourier transform.

#### Beta burst extraction

The beta bursting activity from the CST and EEG were extracted based on previously developed methods, which first calculate the signal’s amplitude within a specific frequency range and then identify instances where amplitude fluctuations surpass a predefined threshold (Little et al., 2019; Shin et al., 2017; Tinkhauser, Pogosyan, Tan, et al., 2017b). Burst analysis was only conducted on CSTs from joint MUs of the 2 grids over the ECU muscle and for trials with at least 5 MUs. Participant specific bursting thresholds were determined by bandpass filtering each trial between 13 to 30 Hz (‘bandpass’ function in MATLAB, zero-phase FIR), taking the upper envelope of the signal (using spline interpolation over local maxima separated by at least 30 samples), and then normalising the signal. The 75^th^ percentile threshold was then calculated for the envelope of each trial within a participant and averaged, to determine the threshold used for that participant. After determining the burst threshold, bursts were defined as periods of longer than 100ms of the signal envelope crossing the threshold, to ensure bursts included at least one full oscillation cycle in the beta range.

#### Time-frequency analysis

The time-frequency analysis of the CST and EEG was performed using the continuous wavelet transform, implemented through MATLAB’s *cwt* function (The MathWorks Inc., 2023) between 1 and 45 Hz. CMC was computed utilizing magnitude-squared wavelet coherence via the *wcoherence* function in MATLAB. A comparable method was applied to track the temporal progression of IMC, employing a custom MATLAB script based on the *wcoherence* function. Similar to the coherence analysis, to assess IMC, the MU pool was divided into two randomly selected subpools of equal size. Magnitude-squared wavelet coherence was then computed between the CSTs of these MU subpools. This process was repeated 100 times, with each iteration using a different MU subpool configuration. The final IMC value was derived by averaging the coherence estimates obtained across all iterations.

Prior to identifying ON and OFF events, EEG, CST and CMC wavelets were z-scored per frequency bins and torque was upsampled to 300 Hz. ON and OFF events were identified in the EEG, CST and CMC wavelets based on the locations of beta bursts identified in the EEG signals. Events of ON bursting were defined as 500ms windows around bursts whose midpoint occurred at least 250 ms from the beginning or end of the 60 second plateau period. ON events were identified for each trial of a participant and then averaged within a trial. To prevent overlap between ON and OFF events, OFF events were defined as the periods between ON events, beginning 250 ms after the previous ON event and ending 250 ms before the next ON event, that were longer than 500 ms. Windows of overlap 50 ms and step size 500 ms were averaged across the OFF periods. ON and OFF events were averaged across trials of participants and percental mismatch was calculated for each participant. The percental mismatch across participants was then averaged. The percental mismatch between ON and OFF events was defined as (ON – OFF)/OFF * 100 and was also calculated for the torque to ensure bursting effects did not result from unstable force production.

### Statistical Analysis

The statistical analysis was conducted using custom MATLAB scripts and the Neurospec 2.11 toolbox (for significant coherences in the IMC and CMC as aforementioned). Results are reported as median and interquartile range (IQR) unless otherwise stated. To analyse significant beta activity differences between ON and OFF events in the time-frequency domain, we applied cluster-based permutation testing based on the approach of Maris and Oostenveld (2007). This method addresses the multiple comparisons problem inherent in multidimensional data (time and frequency) by evaluating clusters of neighbouring samples under a single permutation distribution, rather than assessing each sample independently. We performed 10,000 permutations with a univariate clustering threshold of 0.05, allowing for statistically robust identification of significant clusters in the beta range. After testing for normality using the Shapiro-Wilk test, non-parametric Friedman tests were applied to look at differences in MU count between muscle groups and MVCs. Post-hoc Dunn-Sidak tests were used for multiple comparisons of significant results.

## Results

### MU extraction across muscle locations and force production

The distribution of MUs across muscles (ECR and ECU) and force levels (10%, 20% and 30% MVC) prior to thresholding for ≥5 MUs can be observed in Figure 2 across 24 participants. The median number of MUs identified per trial for both muscles across MVCs (10, 20, 30%), respectively, were 2 (1-4), 1 (0-3) and 1 (0-2) for the ECR, and 4 (2-8), 3 (1-7), and 2 (0-9), for the ECU. A significant between-group difference was observed between the ECU and ECR across all force levels (Friedman test: p_10%_ = 0.0067, p_20%_ = 0.0011, p_30%_ = 0.0004), indicating significantly more MUs in the ECU across all force levels. MU counts in the ECU decreased with increasing MVC, with significantly less MUs in 30% MVC versus 10% MVC (Friedman test: p= 0.00013) and versus 20% MVC (Friedman test: p = 0.021). ECR MU counts also differed significantly between 10% MVC and 30% MVC (Friedman test: p = 0.000082).

**Figure 2.**
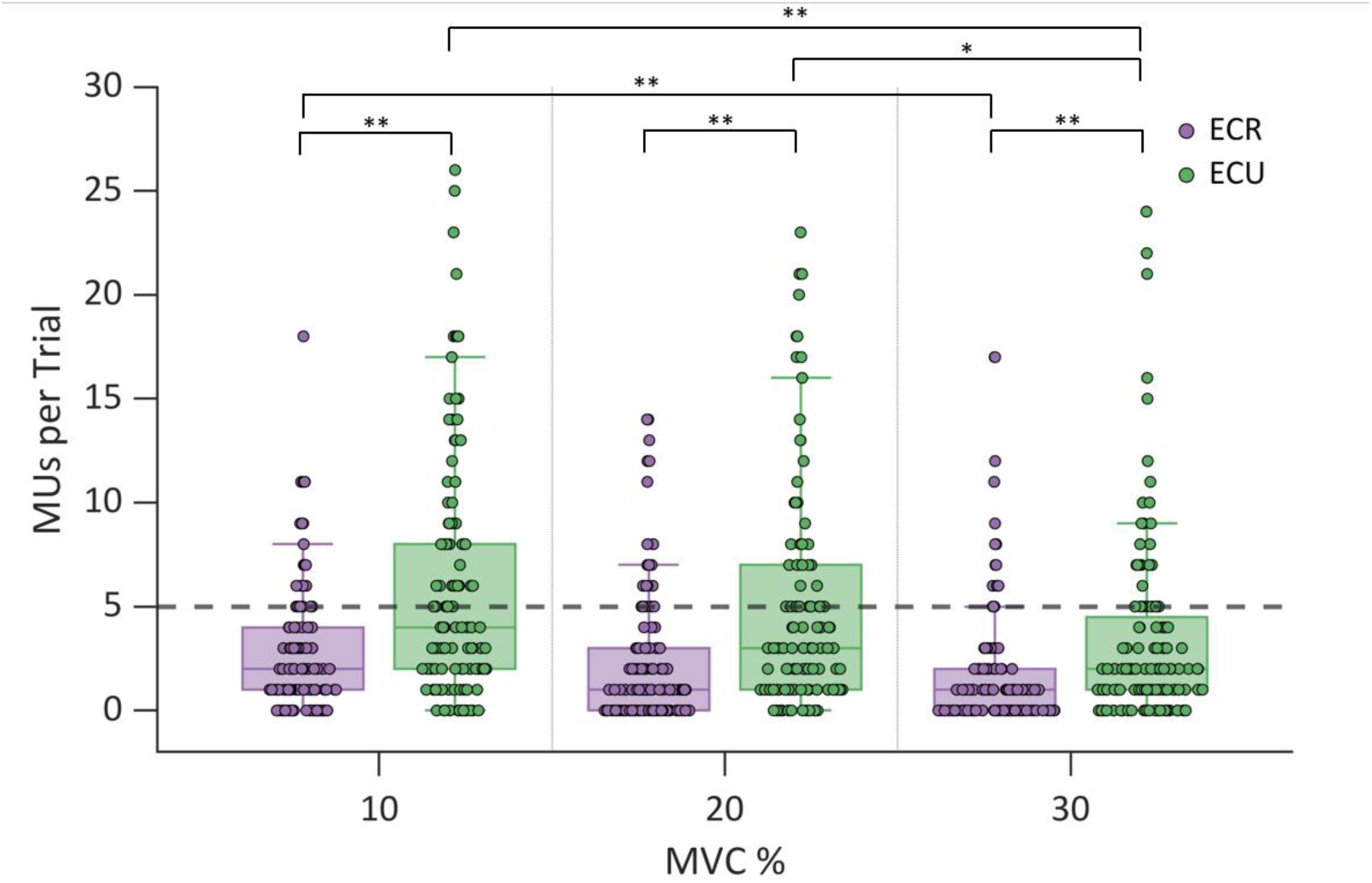
Number of motor units (MUs) per trial identified in extensor muscles for various grid combinations (ECR: extensor carpi radialis proximal (ECRP) + extensor carpi radialis distal (ECRD); ECU: extensor carpi ulnaris proximal (ECUP) + extensor carpi ulnaris distal (ECUD)) and target forces (10%, 20% and 30% maximum voluntary contraction (MVC)). Each scatter marker represents the number of MUs per trials per participant (total trials per muscle = 360, participants=24). * 0.01 < p-value < 0.05; ** 0.001 < p-value < 0.01.

Across all trials with at least one successfully decomposed MU, the median number of MUs per trial per grid across all participants was 2 (1-4). MUs were initially identified from four electrode grids, ECRD, ECRP, ECUD and ECUP, placed over the distal and proximal regions of the two muscles. However, due to low MU yield per individual grid (with only 12.14%, 16.34%, 22.13%, and 30.92% of trials exceeding the ≥5 MU threshold for ECRD, ECRP, ECUD, and ECUP, respectively), data from grids over the same muscle were combined to increase sampling density. This merging yielded a substantially higher proportion of usable trials: 26.69% of ECR trials and 42.21% of ECU trials exceeded the 5 MU threshold after grid-level merging.

Trials with a minimum number of MUs (≥5 MUs) and resampled CSTs across MVCs were included in the coherence analysis, with trial count, participants and MU count across grid configurations seen in Table 1. The MUs extracted demonstrate a decreasing number of MUs with target force, which aligns with previous findings (Del Vecchio, Negro, et al., 2019).

**Table 1.** Motor unit distribution and characteristics by MVC level and grid configuration. Summary of identified motor units (MUs) across individual levels of maximum voluntary contraction (10%, 20%, and 30% MVC) following filtering for grid configurations containing ≥5 MUs. Values are averaged across trials and reported as mean ± SD. ECRD (extensor carpi radialis distal); ECRP (extensor carpi radialis proximal); ECUD (extensor carpi ulnaris distal); ECUP (extensor carpi ulnaris proximal); ECU = ECUD+ECUP; ECR = ECRD+ECRP; ECD = ECRD+ECUD; ECP=ECRP+ECUP; All Grirds = ECRD+ECRP+ECUD+ECUP.

| Muscle | MVC (%) | Participants | Participants Resampled | Trials | Trials Resampled | MUs | MUs Resampled | Discharge Rate | Discharge Rate Resampled |
| --- | --- | --- | --- | --- | --- | --- | --- | --- | --- |
| ECRD | 10 | 5 | 5 | 10 | 10 | 59 | 50 | 14.86±2.66 | 14.73±2.71 |
| | 20 | 3 | 3 | 8 | 8 | 56 | 40 | 13.68±2.19 | 13.74±2.45 |
| | 30 | 2 | 2 | 3 | 2 | 22 | 10 | 15.31±2.25 | 15.13±2.40 |
| ECRP | 10 | 3 | 2 | 10 | 8 | 58 | 40 | 12.89±1.11 | 13.26±1.50 |
| | 20 | 3 | 3 | 7 | 7 | 53 | 35 | 14.03±2.46 | 14.04±2.48 |
| | 30 | 3 | 2 | 8 | 6 | 63 | 31 | 14.20±1.52 | 14.99±0.60 |
| ECUD | 10 | 5 | 5 | 21 | 21 | 155 | 116 | 13.82±1.00 | 14.01±0.97 |
| | 20 | 6 | 6 | 19 | 16 | 161 | 89 | 15.17±1.02 | 15.34±1.12 |
| | 30 | 4 | 3 | 12 | 7 | 97 | 40 | 16.22±1.26 | 16.45±1.75 |
| ECUP | 10 | 10 | 10 | 35 | 35 | 302 | 232 | 14.00±1.29 | 14.10±1.34 |
| | 20 | 9 | 9 | 27 | 26 | 207 | 152 | 15.06±1.23 | 15.17±1.30 |
| | 30 | 4 | 4 | 15 | 14 | 117 | 72 | 16.08±1.29 | 16.32±1.52 |
| ECU | 10 | 15 | 15 | 55 | 51 | 594 | 342 | 13.91±1.31 | 14.10±1.40 |
| | 20 | 14 | 14 | 45 | 43 | 468 | 271 | 14.95±1.34 | 15.12±1.53 |
| | 30 | 11 | 11 | 30 | 30 | 277 | 193 | 15.72±1.57 | 16.02±1.67 |
| ECR | 10 | 9 | 8 | 25 | 23 | 193 | 133 | 13.86±2.53 | 14.28±2.77 |
| | 20 | 7 | 7 | 23 | 21 | 183 | 124 | 13.30±1.67 | 13.35±1.94 |
| | 30 | 4 | 3 | 15 | 9 | 129 | 56 | 14.18±1.38 | 15.12±2.40 |
| ECD | 10 | 13 | 13 | 38 | 38 | 315 | 222 | 14.14±1.77 | 14.49±1.77 |
| | 20 | 10 | 10 | 31 | 30 | 292 | 170 | 14.53±1.42 | 14.96±1.59 |
| | 30 | 8 | 8 | 20 | 15 | 171 | 94 | 15.53±1.51 | 15.76±1.78 |
| ECP | 10 | 13 | 13 | 46 | 46 | 455 | 315 | 13.74±1.25 | 13.89±1.31 |
| | 20 | 13 | 13 | 34 | 31 | 323 | 206 | 14.64±1.48 | 14.85±1.54 |
| | 30 | 5 | 5 | 17 | 16 | 193 | 105 | 15.34±1.75 | 15.73±1.93 |
| All Grids | 10 | 21 | 21 | 73 | 71 | 927 | 514 | 14.07±1.79 | 14.45±2.01 |
| | 20 | 19 | 18 | 63 | 59 | 764 | 434 | 14.69±1.71 | 15.10±1.85 |
| | 30 | 13 | 13 | 40 | 37 | 475 | 307 | 15.15±1.81 | 15.53±1.90 |

### Intramuscular coherence

IMC across different grid combinations was investigated for the shared neural drive of the extensor muscles, which act synergistically during wrist extensions (Figure 3). This analysis aims to examine the functional coordination both within and between these muscles, as IMC indicates whether muscles work together as part of a functionally related group or a neural synergy (Laine & Valero-Cuevas, 2017). Understanding these coherence patterns can help identify which extensor muscle regions exhibit enhanced beta band frequencies. While all force levels and grid combinations show significant alpha (8-13 Hz) coherence, stronger and significant coherences in the beta band was observed consistently across MVCs only in the grids placed over the distal and proximal regions of the ECU (ECUD and ECUP), as well as in the shared coherence between these two grids. This suggests that specific MUs, where beta rhythms are more dominant or synchronised, are primarily located in the ECU. These findings provide insight into where and how beta activity manifest in upper limb muscle activity. The limit for defining significant coherences was 0.034, corresponding to the 95% confidence limit determined by the number of segments in the Neurospec analysis. Furthermore, the ECU beta common neural drive increased with force with a non-significant peak of 0.033 at 22.5Hz at 10% MVC, a significant peak of 0.037 at 21Hz at 20% MVC and a significant peak of 0.048 at 24.5Hz at 30% MVC. These findings show that, similar to (Hug et al., 2021), muscles of the same synergistic group do not necessarily share the same neural drive, as the ECR and ECU share minimal common drive in this case. Despite increasing the number of MUs in the calculations when merging ECR and ECU, the IMC does not increase, further emphasizing this lack of shared neural drive.

**Figure 3.**
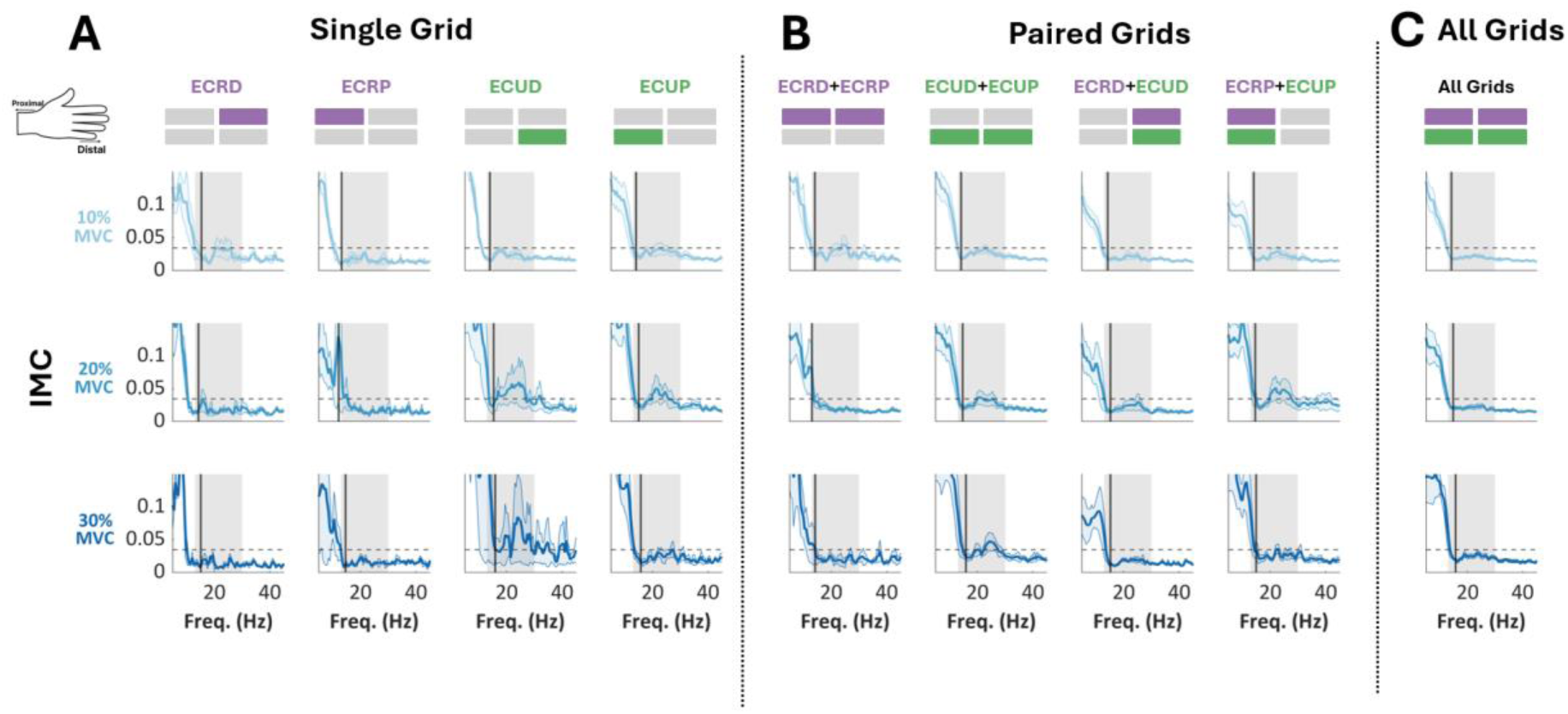
Intramuscular coherences across extensor muscles and force levels (10, 20, 30% MVC). Horizontal dashed line is 95% significant threshold. Vertical solid line is mean discharge rate. Shaded area is standard error of mean. ECRD (extensor carpi radialis distal); ECRP (extensor carpi radialis proximal); ECUD (extensor carpi ulnaris distal); ECUP (extensor carpi ulnaris proximal); All Grids = ECRD+ECRP+ECUD+ECUP.

### Corticomuscular coherence

The corticomuscular coherence (CMC) between cortical (EEG electrode C3 signals) and muscle (common input to decomposed ECU MUs quantified by the CST), was estimated across three force levels (10%, 20% and 30% MVC). The coherence spectrum can be seen in Figure 4. At 10% MVC, coherence remained relatively low across all frequencies, with no strong beta-band peak observed. At 20% MVC, coherence was slightly higher but still exhibited significant beta-band activity in the ECUD grid (0.032 at 21.09Hz). At 30% MVC, coherence increased noticeably, particularly in the beta range, where a significant peak was present at 25.2Hz (coherence 0.05), and 22.27Hz (coherence 0.048) in the ECRD and ECUD grids respectively. These results show that beta band coherence becomes more pronounced with increasing force production in the distal forearm, supporting the idea that the synchronization between the motor cortex (C3) and ECU MUs strengthens as contraction levels rise. This suggests that corticospinal drive in the beta range plays an increasing role in muscle activation at higher force levels.

**Figure 4.**
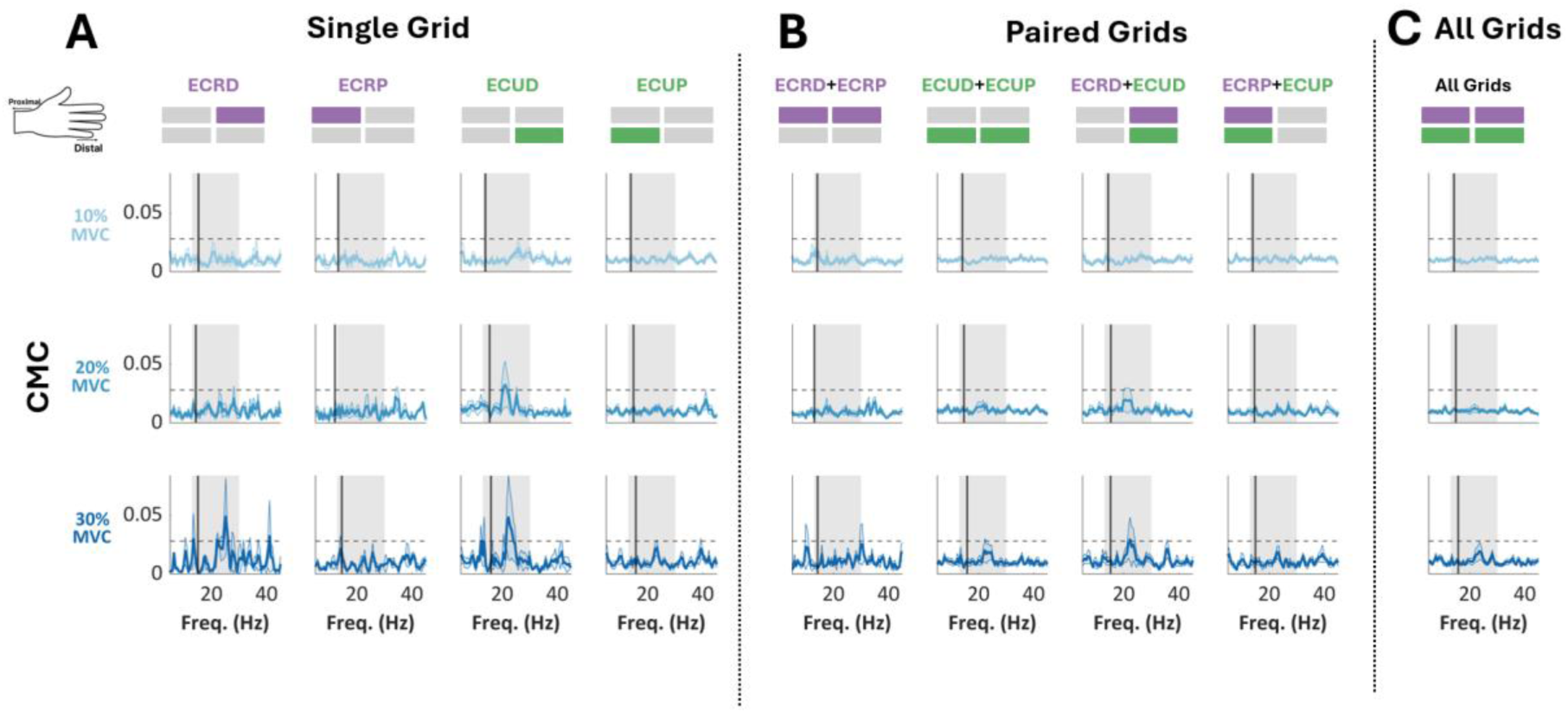
Corticomuscular coherence –. Average CMC (solid line) between ECU CST and EEG C3 averaged across trials and participants. Standard error mean (SEM) indicated by shaded area. 95% confidence limit (dashed line) and mean discharge rate (vertical solid line). ECRD (extensor carpi radialis distal); ECRP (extensor carpi radialis proximal); ECUD (extensor carpi ulnaris distal); ECUP (extensor carpi ulnaris proximal); All Grids = ECRD+ECRP+ECUD+ECUP.

Figure 5 presents data combined across all MVC levels and highlights the rationale for selecting the ECU muscles as the primary focus for burst analysis. Among all electrode configurations, only the ECU demonstrated a consistent and statistically significant IMC beta-band peak, alongside the highest number of contributing participants (n=20). In contrast, CMC beta activity was only observed in the ECUD and did not reach statistical significance. The raw (un-resampled) MU counts, trials, and participants contributing to Figure 5 are detailed in Table 2.

**Figure 5.**
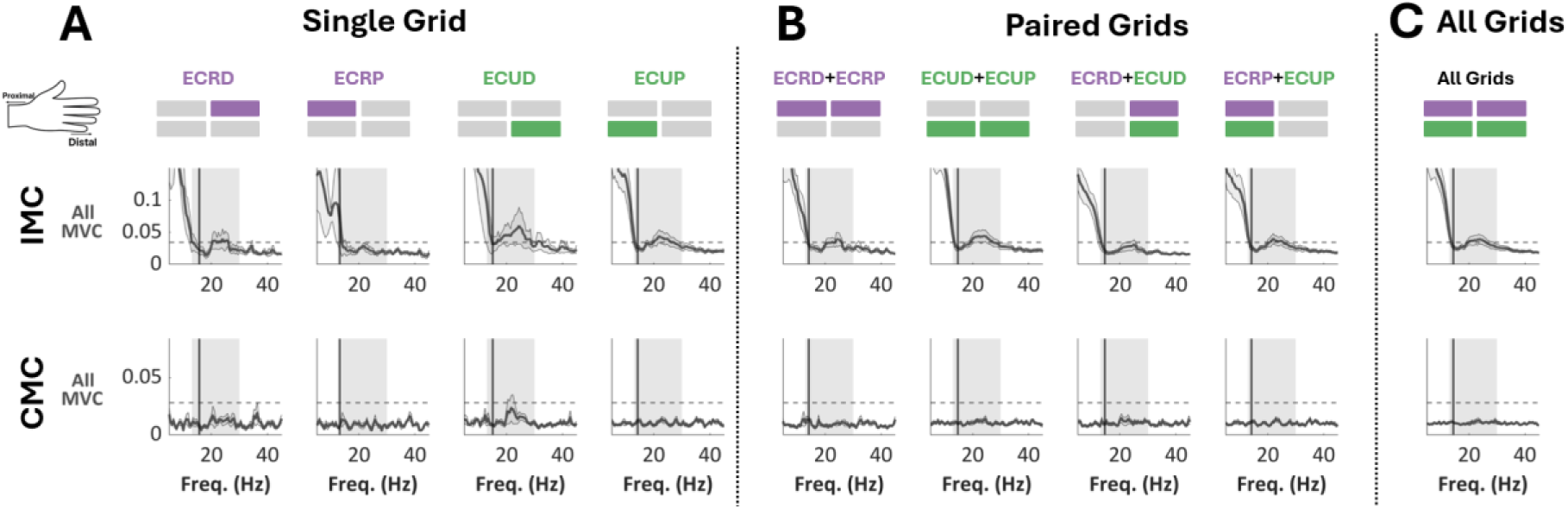
Intramuscular and corticomuscular coherence across extensor muscles at all force levels together (10, 20, 30% MVC). CSTs have not been resampled for consistent firings. ECRD (extensor carpi radialis distal); ECRP (extensor carpi radialis proximal); ECUD (extensor carpi ulnaris distal); ECUP (extensor carpi ulnaris proximal); All Grids = ECRD+ECRP+ECUD+ECUP.

**Table 2.** Motor unit distribution and characteristics across grid configurations (all MVC levels combined). Summary of identified motor units (MUs) across all MVC levels (10%, 20%, and 30% MVC) merged into a single analysis, filtered for ≥5 MUs per configuration. Values are averaged across trials and reported as mean ± SD. Muscle grid abbreviations as in Table 1.

| Muscle | Participants | Trials | MUs | Discharge Rate |
| --- | --- | --- | --- | --- |
| ECRD | 5 | 21 | 137 | 14.48±2.41 |
| ECRP | 4 | 25 | 174 | 13.63±1.74 |
| ECUD | 7 | 52 | 413 | 14.87±1.42 |
| ECUP | 12 | 77 | 626 | 14.78±1.49 |
| **ECU** | **20** | **130** | **1339** | **14.69±1.56** |
| ECR | 10 | 63 | 505 | 13.73±2.00 |
| ECD | 14 | 89 | 778 | 14.59±1.67 |
| ECP | 15 | 97 | 971 | 14.33±1.54 |
| **All Grids** | **22** | **176** | **2166** | **14.53±1.81** |

Focusing specifically on the ECU muscle, 130 trials from 20 participants—representing 36.1% of all trials and 83.3% of participants—were included in the burst analysis, combining trials across all force levels (10%, 20%, and 30% MVC) from the joint ECU grids. This selection was based on the observation that grids covering the ECU yielded a greater number of identified MUs and stronger common beta drive compared to those covering the ECR.

Beta activity, though transient, is evident across brain, muscles, and their connectivity during sustained extensor contraction. In Figure 6, time periods in the CST, CMC, and torque have been identified based on the ‘ON’ and ‘OFF’ periods of the EEG beta bursts. When averaging across trials, beta ON events appear brief and clearly distinguishable from OFF periods, not only in the EEG but also in the CST and CMC. This emphasizes the transient nature of beta bursts beyond the cortex. Cluster-based permutation analyses confirmed significant transient differences between ON and OFF periods in both muscle beta power within the CST and brain-muscle connectivity in the CMC. Importantly, no significant modulation in grip force was observed during beta ON events, with force fluctuations remaining within 1% between ON and OFF periods, indicating that the observed beta bursts were not driven by changes in motor output. These bursts were synchronized across cortical and peripheral sites, with CST beta activity occurring simultaneously to EEG bursts.

**Figure 6.**
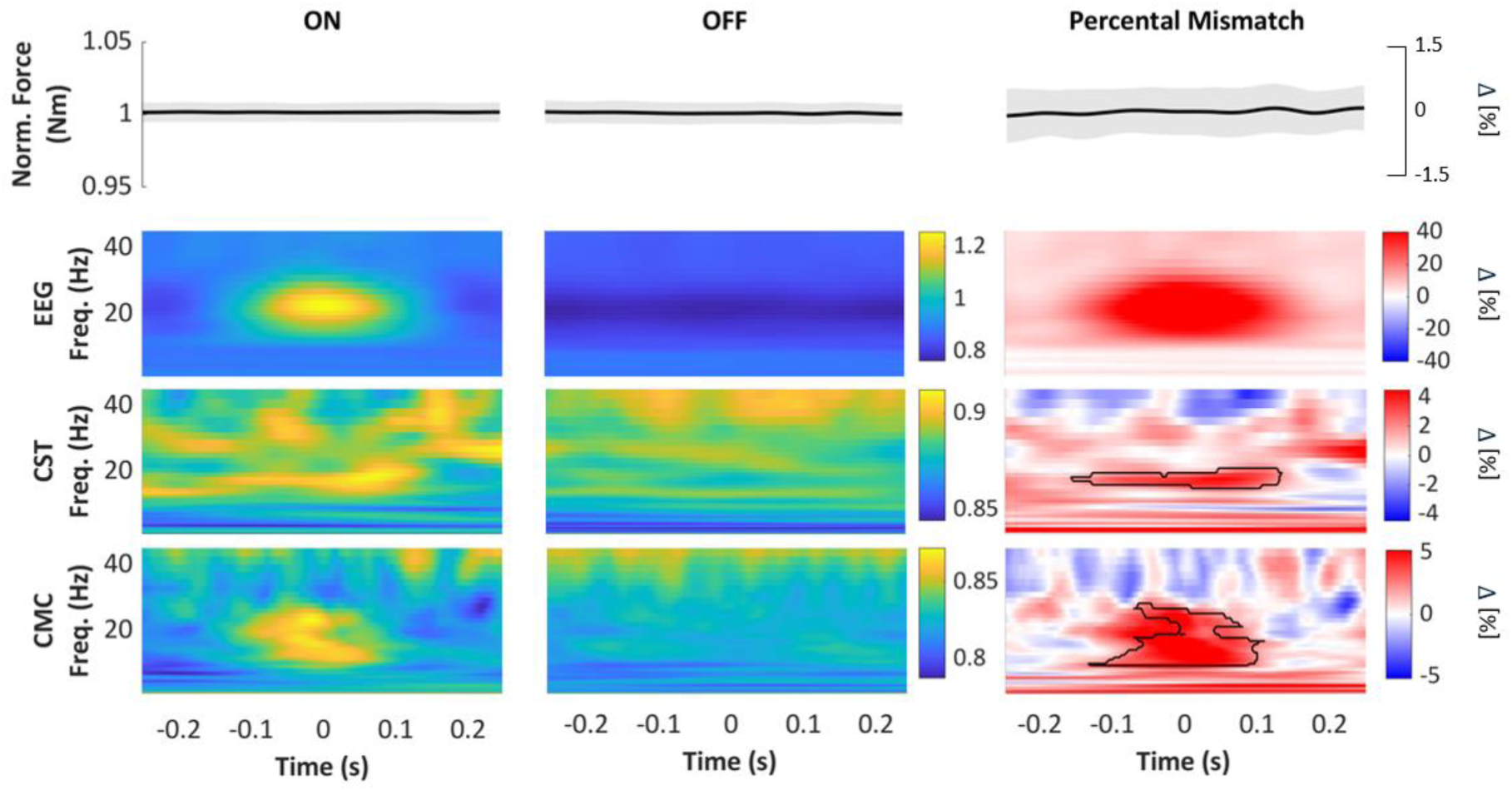
Neural and peripheral bursting activity at burst occurrences in the EEG. Burst ON and OFF events were averaged across trials. From top to bottom: normalised force, wavelet transformed EEG, CST and CMC with t=0 representing the centre of ON periods (left), OFF periods (centre) and percental mismatch between ON and OFF periods (right). Black outlines represent the most significant clusters in the beta band (p<0.05).

## Discussion

We investigated the cortical-peripheral relationship of beta activity and bursts between the output of spinal motor neurons and neural oscillations during isometric forearm contractions, providing novel insights into how beta activity is transmitted across the motor system in the upper limb. We assessed the reliability of HDsEMG for detecting beta bursts across various force levels and extensor muscle regions and examined the alignment of beta bursts at the cortical and peripheral levels. Our findings reveal that beta activity in MU pools of the upper limb occurs in discrete, transient bursts that strongly reflect and overlap with cortical beta bursts, extending previous observations made in the lower limb (Bräcklein et al., 2022). We also show that the ECU exhibits stronger shared beta neural input than the ECR, highlighting muscle-specific differences in common beta input. Similar to previous corticomuscular beta burst research (Bräcklein et al., 2022; Echeverria-Altuna et al., 2022; Simpson et al., 2024), CST beta bursts occurred simultaneously with EEG beta bursts. These findings provide new insights into corticospinal dynamics during voluntary muscle contractions by being the first to examine beta burst synchronization between the cortex and MUs in the upper limb, and by exploring how these bursts vary across different force levels.

## Differences in shared neural drive to extensor muscles

Our results suggest that extensor muscles within the same functional group, namely the ECU and ECR, differ in the strength of shared beta-band neural input, as evidenced by significantly high beta synchronisation across all MVCs in the ECU, but not ECR. This indicates that the distribution of common beta input is not uniform across forearm extensors and may reflect muscle-specific modulation of synaptic drive within the beta range. While we cannot determine from this data whether the sources of input differ qualitatively between muscles, the disparity in synchronisation strength may relate to differences in functional roles and control strategy. Whilst the ECU primarily stabilises the wrist and facilitates both wrist extension and ulnar deviation (movement of the wrist toward the little finger) (Finsen et al., 2005), the ECR provides a balancing force during wrist extension, counteracting excessive ulnar deviation and contributing to overall wrist extension (Sawyer et al., 2023). Given their biomechanical distinctions, the two muscles may activate independently based on specific movement demands, potentially requiring greater synchronisation among MUs in the ECU to support its stabilising role. While functionally related muscles often exhibit a stronger shared drive (Kerkman et al., 2018; Laine et al., 2015), our findings imply that this relationship may vary according to each muscle’s biomechanical function. Finally, exploratory analyses did not reveal consistent inter-muscle beta burst synchrony between ECU and ECR, suggesting that coordinated beta transmission across these synergists may be weak or variable during low-force isometric tasks.

The presence of alpha band (8-12 Hz) IMC across all muscles may reflect synchronized oscillations associated with motor control and stability, as previously observed in both upper and lower limb tasks (Boonstra et al., 2008, 2015; de Vries et al., 2016). This low-frequency coupling may relate to the minimum discharge rate required for MU recruitment and could serve to enhance postural stability or regulate movement execution, particularly during slower, controlled contractions (Danna-Dos-Santos et al., 2014; Kouzaki & Masani, 2012). However, in this study, we assumed that activity from the ECR and ECU could be distinguished in HDsEMG recordings, despite potential signal contamination from coactivation of surrounding extensor muscles, such as the extensor digitorum, and inter-individual variations in muscle anatomy relative to grid placements. While minor differences were observed between distal and proximal grids, they were grouped together due to the limited number of decomposed MUs. As expected from previous studies on the upper limb (Rojas-Martínez et al., 2012, 2020), we observed a low number of decomposed MUs from the extensors, likely due to the fusiform architecture of the ECU and ECR. Compared to pennate muscles, such as the tibialis anterior, fusiform muscles have parallel muscle fibres and exhibit less variety in MU action potential shapes towards the skin surface, which may reduce the effectiveness of decomposition techniques (Francic & Holobar, 2021).

## Corticomuscular beta transmission and relationship to force

We observed a trend toward increasing beta CMC with higher force levels; however, statistical significance was not fully achieved, likely due to inter-individual variability (Fisher et al., 2012; Ushiyama et al., 2011). This trend aligns with the notion that stronger corticospinal coupling may support forceful contractions, as previously shown in muscles such as the first dorsal interosseous, tibialis anterior and knee flexors/extensors at low-to-moderate forces (Dal Maso et al., 2017; McManus et al., 2019; Ushiyama et al., 2012). However, findings across the literature remain mixed, with some reporting a decreasing MU synchronisation strength at higher forces (Kline & De Luca, 2015) and others showing no clear trend(Christou et al., 2007), likely due to a limitation of MU discrimination at lower contractions intensities.

Our findings are further contextualized by recent evidence from Abbagnano et al. (2025), who showed that cortical beta band oscillations are uniformly projected to the entire motor neuron pool across a full range of recruitment thresholds in the tibialis anterior. In their study, CMC remained stable across contraction intensities, and conduction delay estimates confirmed consistent corticospinal transmission via fast-conducting fibres, regardless of MU size. While our results show a non-significant trend toward increased beta CMC with force, the lower overall coherence values and inter-individual variability observed in our upper-limb dataset may reflect anatomical or functional differences from the lower limb. Unlike the tibialis anterior, a unipennate muscle with a relatively homogeneous MU pool, the ECR and ECU are smaller, fusiform muscles that may receive more distributed or muscle-specific descending inputs. Taken together, these findings suggest that while the core transmission mechanism for beta input may be conserved across muscles, its observable strength and variability could be influenced by muscle architecture and task demands.

## Reliability of HDsEMG in detecting beta bursts in the upper limb

Our findings confirm that HDsEMG reliably detects transient beta bursts in forearm extensor muscles, reinforcing its utility for studying upper limb corticospinal dynamics. We found that peripheral beta bursts were not only transient, but synchronized with cortical bursts, with CST beta activity being time-locked to EEG bursts, similar to findings by Bräcklein et al. (2022) and Echeverria-Altuna et al. (2022). Since no significant changes in produced torque were observed in relation to the timing of beta bursts, it is unlikely that these bursts play a direct role in force modulation. Yet, Zicher et al. (2023) found that when beta activity was modelled as transient bursts rather than continuous oscillations, these bursts reached levels capable of inducing small fluctuations in force, with estimated common beta inputs to motor neurons aligning with experimental observations. Similarly, Echeverria-Altuna et al. (2022) demonstrated that brain-muscle neural synchronisation occurs in brief, frequency-specific bursts, even during sustained motor tasks, further supported by our results. Beta bursts may play an indirect role in sustaining motor activity, potentially by monitoring peripheral states (Baker, 2007; Witham et al., 2011b) or through alternative neural pathways not examined in this study, such as intracortical connectivity or non-linear cortical-peripheral integration of sensory information. However, further research is required to confirm these hypotheses.

## Limitations

Despite the strengths of this study, several methodological limitations should be considered when interpreting the findings. First, given the target muscle size and limitations inherent with non-invasive methods, such as HDsEMG, the ability to detect deep MUs accurately in the forearm muscles or a high number of decomposed MUs was hindered. Second, trials with low MU counts were discarded, which could introduce sampling bias and affect the generalizability of the findings. Lastly, the variability in force maximum and muscle size across participants further complicates the interpretation, as strength and beta power were seemingly correlated.

## Conclusion

Future research should investigate how corticomuscular beta bursting behaves across a range of task conditions, including dynamic movements and prolonged exertion, to better understand how these bursts contribute to motor control during more natural, functional tasks and provide further ecological validity to the observed beta burst dynamics. Extending these findings to clinical populations, such as individuals with Parkinson’s disease, stroke, or those using prosthetic devices, could yield crucial information on how altered corticospinal coupling impacts motor behaviour in these populations. To ensure broader generalisability, future studies should examine other muscles and incorporate intracortical recordings or high-density EEG to refine coherence analysis.

In conclusion, this study provides novel insights into how beta bursts contribute to corticospinal interactions during motor control, specifically in the upper limb muscles and across force levels. Our findings advance our understanding of the neural drive and motor stability in reflecting cortical beta activity, as well as understanding the time-locking of cortical and peripheral bursts. These insights could have important implications for neurorehabilitation strategies and prosthetic control, particularly for individuals with motor impairments, such as Parkinson’s disease, where beta bursts are a well-established neural biomarker of its motor symptoms, particularly rigidity and bradykinesia. Moreover, the ability to reliably detect peripheral beta bursting may inform the development of closed-loop DBS systems by providing an accessible biomarker for adaptive neuromodulation. Tracking beta dynamics at the MU level could allow more precise, feedback-driven stimulation paradigms targeting abnormal beta synchronization in Parkinson’s disease. By validating the ability to track beta bursts across the corticospinal tract in upper limb muscles, similar to the tibialis anterior, using HDsEMG as a robust neural signal, we pave the way for future innovations in neuroengineering, particularly for investigating changes in the brain during neuromodulation interventions.

## Acknowledgements

CG was supported by UK Research and Innovation [UKRI Centre for Doctoral Training in AI for Healthcare grant number EP/S023283/1]. APV was supported by the European Union’s Horizon Europe research and innovation programme under the Marie Skłodowska-Curie grant agreement No 101151398.

## Notes

### Competing Interest Statement

The authors have declared no competing interest.

